# Identifying a task-invariant cognitive reserve network using task potency

**DOI:** 10.1101/747311

**Authors:** A.C. van Loenhoud, C. Habeck, W.M. van der Flier, R. Ossenkoppele, Y. Stern

## Abstract

Cognitive reserve (CR) is thought to protect against the consequence of age- or disease-related structural brain changes across multiple cognitive domains. The neural basis of CR may therefore comprise a functional network that is actively involved in many different cognitive processes. To investigate the existence of such a “task-invariant” CR network, we measured functional connectivity in a cognitively normal sample between 20-80 years old (N=265), both at rest and during the performance of 11 separate tasks that aim to capture four latent cognitive abilities (i.e. vocabulary, episodic memory, processing speed, and fluid reasoning). For each individual, we determined the change in functional connectivity from the resting state to each task state, which is referred to as “task potency” (Chauvin et al., 2017; Chauvin et al., 2018). Task potency was calculated for each pair among 264 nodes (Power et al., 2011) and then summarized across tasks reflecting the same cognitive ability. Subsequently, we established the correlation between task potency and premorbid IQ or education (i.e. CR factors). We identified a set of 57 pairs in which task potency showed significant correlations with IQ, but not education, across all four cognitive abilities. These pairs were included in a principal component analysis, from which we extracted the first component to obtain a latent variable reflecting task potency in this task-invariant CR network. This task potency variable moderated the relationship between cortical thickness and episodic memory performance (β=−.64, p=.01), and showed a direct effect on fluid reasoning (β=.08, p<.01) after adjusting for the effects of cortical thickness. Our identification of this task-invariant network contributes to a better understanding of the mechanism underlying CR, which may facilitate the development of CR-enhancing treatments. Our work also offers a useful alternative operational measure of CR future studies.

## Introduction

Cognitive reserve (CR) describes the ability to maintain cognitive function in the presence of age- or disease-related structural brain changes.^1^ Individuals with higher educational attainment, premorbid IQ, and physical or cognitive activity, among other factors, generally have greater CR.^2–10^ CR is often conceptualized as a moderator between brain changes and cognition, such that the cognitive impact of these changes is attenuated at higher levels of CR. Another common way to demonstrate CR is the presence of a direct positive relationship with cognition after adjusting for brain status.^11^ These effects of CR occur across various cognitive domains, and therefore the neural basis of CR may comprise a functional network that is actively involved in many different cognitive processes. Functional magnetic resonance imaging (fMRI) has provided evidence for the existence of such a “task-invariant” mechanism underlying CR. Based on the comparison of BOLD activation patterns from multiple cognitive tasks, several studies identified common regions of activity across conditions.^12–14^ Recruitment of these regions showed direct associations with CR factors, such as education^12^ and premorbid IQ.^13,14^ In addition, resting state fMRI, which has the advantage of providing information about the brain’s functional organization rather than regional activity, has also been used in the context of CR. These studies have suggested that CR is represented by global connectivity of specific “cognitive control” regions,^15–17^ which support cognition at a task-invariant level by mediating flexible adaptation to changing task demands.

Although there is general consensus on the existence of a link between resting state functional connectivity and cognition,^18^ this link is indirect in nature as these fMRI data are not acquired during task performance. In fact, several studies have shown that functional connectivity is dynamic,^19,20^ and systematic differences between rest and task states exist despite overall preservation of network topography.^21,22^ This is consistent with the idea that neural responses evoked by tasks build upon an already present functional connectivity baseline.^23,25^ These systematic differences between resting state and task-based functional connectivity have motivated the development of novel fMRI analysis techniques, such as the “task potency” method.^26–27^ Task potency captures a brain region’s functional connectivity during task performance after adjusting for its resting state baseline and thus reflects connectivity changes that occur in response to an experimental condition. As the task potency method provides task-related information while also taking into account the interconnected nature of the brain, it offers an ideal approach for the identification of a network that is actively involved in multiple tasks.^27^

In the present study, we used a technique that resembles the task potency method to investigate the existence of a task-invariant CR network. We used data from the Reference Ability Neural Network (RANN) study,^28^ in which healthy individuals across the age span (i.e. 20-80) underwent fMRI during rest and while performing a large set of cognitive tasks. These tasks reflect four latent cognitive abilities that capture most of the age-related variance in cognitive performance: vocabulary, episodic memory, processing speed, and fluid reasoning.^29–31^ Based on 11 RANN tasks, we calculated task potency maps for each cognitive ability and then assessed the relationship with known CR factors (i.e. education and premorbid IQ). We aimed to find a common network of connections in which task potency consistently correlated with these CR factors across all cognitive abilities. To test if the network behaved in accordance with the CR theory, we established its influence on cognition relative to the effects of brain structure. We predicted that greater task potency in this network (which would suggest greater involvement during tasks) was associated with either an attenuated impact of lower cortical thickness on cognition, or an improved cognitive performance after adjusting for cortical thickness.

## Material and methods

### Participants

All subjects participated in the RANN study; they were recruited through random market mailing and provided informed consent prior to participation. Subjects were required to be aged between 20-80 years, native English speakers, right-handed, and have at least a fourth grade reading level. Subjects were screened for MRI contraindications and hearing or visual impairment that would impede testing. Subjects were free of medical or psychiatric conditions that could affect cognition. Careful screening ensured that the elder subjects did not meet criteria for dementia or Mild Cognitive Impairment (MCI). A score greater than 130 was required on the Mattis Dementia Rating Scale^32^ to ensure cognitive normalcy. Further, participants were required to have no or minimal complaints on a functional impairment questionnaire.^33^ From this RANN cohort, we selected all subjects (n=323) who completed fMRI scans during rest and while performing 11 RANN tasks (as described below, Picture Naming was not considered for analysis here). Our final sample consisted of 265 subjects, after exclusion of those with high proportions of missing or scrubbed fMRI data (Figure 1). Individuals who were excluded had a higher mean age and lower global mean cortical thickness; there were no other differences compared to the included sample (Supplementary Table 1).

**Figure 1.**
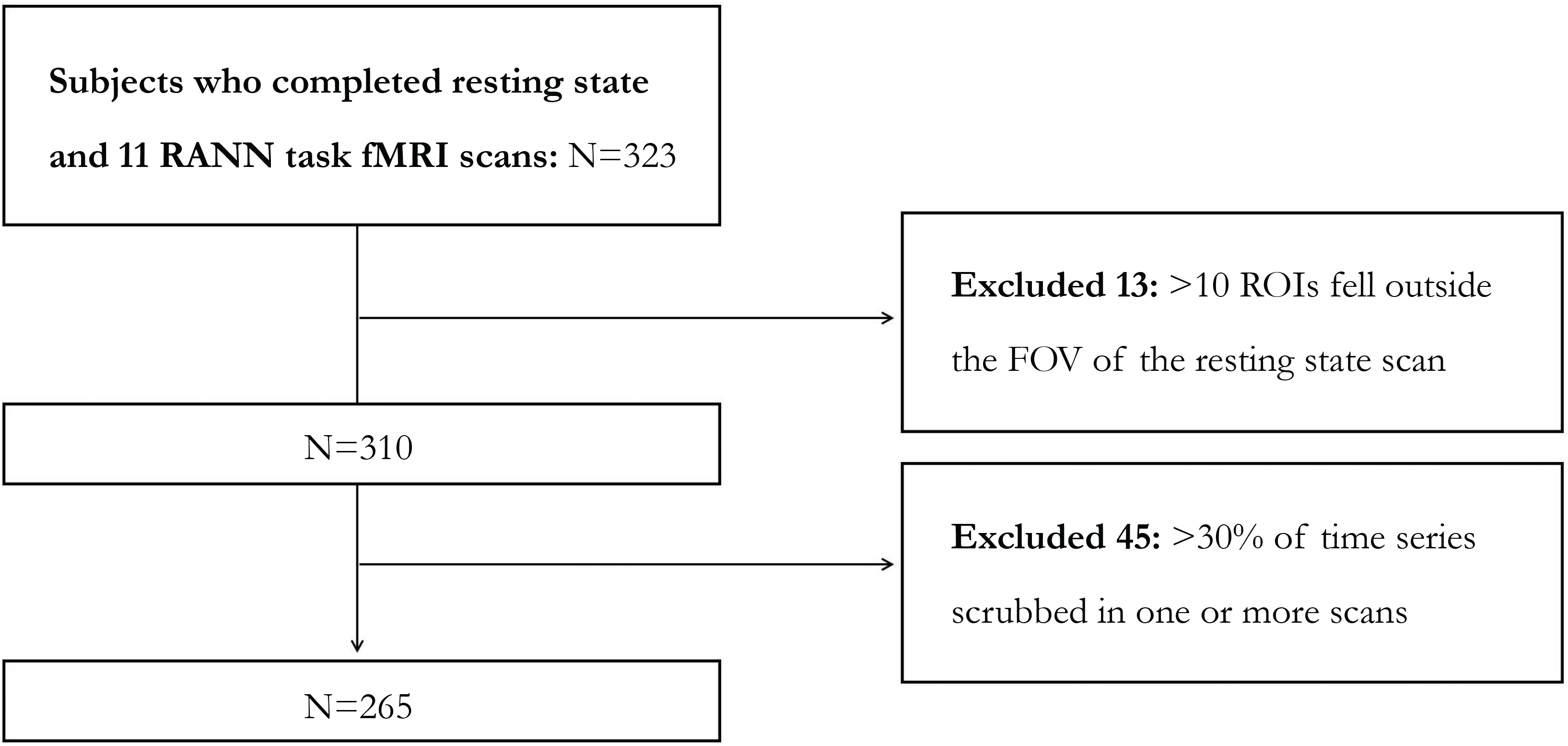
Flow chart illustrating the inclusion/exclusion of individuals in this study. ROI=region of interest according to Power et al (2011), FOV=field of view.]

### Cognitive reserve factors

We measured education and premorbid IQ as they are both contributing factors of CR.^2^ Education was measured in years, and premorbid IQ was estimated based on American National Adult Reading Test (NART) scores.^34^

### Cognitive assessment

Task stimuli were back-projected onto a screen located at the foot of the MRI bed using an LCD projector. Participants viewed the screen via a mirror system located in the head coil and, if needed, had vision corrected to normal using MR compatible glasses (manufactured by SafeVision, LLC. Webster Groves, MO). Responses were made on a LUMItouch response system (Photon Control Company). Task administration and collection of reaction time (RT) and accuracy data were controlled by EPrime running on a PC. Task onset was electronically synchronized with the MRI acquisition computer.^28^

We used all tasks from the RANN study, except one (i.e. Picture Naming). Since this task requires verbal responses in the scanner, it presumably evokes functional connectivity patterns related to speech production that are not present in other tasks. Moreover, verbally responding considerably increased the amount of head movement during the scan, which may introduce task-related biases and thus further complicates the comparison with other RANN tasks. For the remaining 11 tasks, functional connectivity was computed based on the entire duration of each scan (e.g. for episodic memory tasks, both the encoding and retrieval conditions were included). We briefly describe each RANN tasks below, grouped by their four latent cognitive abilities, more details are provided elsewhere.^28^ Scores on each task were standardized using the total baseline sample of the RANN study as a reference group (N=396), and for each cognitive ability we created a summary score by averaging z-scores from tasks within the same ability.

#### Vocabulary (VOCAB)

The primary dependent variable for all VOCAB tasks is the proportion of correct items. <u>Antonyms</u>:^35^ participants match a given word to its antonym, or to the word most different in meaning. <u>Synonyms</u>:^35^ subjects have to match a given word to its synonym or to the word most similar in meaning, by selecting one option from a set of other words. In both cases, the probe word is presented in all capital letters at the top of the screen, and four numbered choices are presented below.

#### Episodic memory (MEM)

The primary dependent variable for the memory tests is proportion of correctly answered questions. <u>Logical Memory</u>: stories are presented on the computer screen. The subject is asked to answer detailed multiple-choice questions about the story, with four possible answer choices. <u>Paired Associates</u>: pairs of words are presented, one at a time, on the screen, and subjects are instructed to remember the pairs. Following the pairs, they were given a probe word at the top of the screen and four additional word choices below. Subjects were asked to choose the word that was originally paired with the probe word. <u>Word Order Recognition</u>: a list of twelve words is presented one at a time on the screen, and subjects are instructed to remember the order in which the words are presented. Following the word list they are given a probe word at the top of the screen, and four additional word choices below. They are instructed to choose out of the four options the word that immediately followed the word given above.

#### Perceptual speed (SPEED)

The primary dependent variable for all SPEED tasks is reaction time. <u>Letter Comparison</u>:^36^ in this task, two strings of letters, each consisting of three to five letters, are presented alongside one another. Subjects indicate whether the strings are the same or different using a differential button press. <u>Pattern Comparison</u>:^36^ two Figures consisting of varying numbers of lines connecting at different angles are presented alongside one another. Subjects indicate whether the Figures were the same or different by a differential button press. <u>Digit Symbol</u>: a code table is presented on the top of the screen, consisting of numbers one through nine, each paired with an associated symbol. Below the code table an individual number/symbol pair is presented. Subjects are asked to indicate whether the individual pair is the same as that in the code table using a differential button press. Subjects are instructed to respond as quickly and accurately as possible.

#### Fluid reasoning (FLUID)

The primary dependent variable for FLUID tasks is the proportion of correct trials completed. <u>Matrix Reasoning (adapted from Raven)</u>:^47^ subjects are given a matrix that is divided into nine cells, in which the Figure in the bottom right cell is missing. Below the matrix, they are given eight Figure choices, and they are instructed to evaluate which of the Figures would best complete the missing cell. <u>Paper Folding</u>:^38^ subjects select a pattern of holes (from five options) that would result from a sequence of folds in a piece of paper, through which a hole is then punched. The sequence is given on the top of the screen, and the five options are given in a row below. Response consisted of pressing one of five buttons corresponding to the chosen solution. <u>Letter Sets</u>:^38^ subjects are presented with five sets of letters, where four out of the five sets have a common rule (i.e. have no vowels), with one of the sets not following this rule. Subjects are instructed to select the unique set.

### MRI acquisition

All MR images were acquired on a 3.0 T Philips Achieva Magnet scanner. Participants underwent two imaging sessions of approximately two hours. Each session started with a scout, T1-weighted image to determine patient position, after which an MPRAGE scan, 11 fMRI tasks, a resting state BOLD (ranging between 5 and 9.5 minutes), and other imaging modalities were obtained (i.e. FLAIR, DTI, and ASL, which will not be used in this study). The MPRAGE parameters were: TR=6.6 ms, TE=3.0 ms, flip angle=8°, FOV=256×256 mm, matrix size=256×256 mm, number of slices=165, voxel size=1×1×1 mm3. The EPI parameters were: TR=2000 ms, TE=20 ms, flip angle=72°, FOV =224×132 mm, matrix size=112×110 mm, number of slices=33, voxel size=2×2×2 mm3 (task scans); TR=2000 ms, TE=20 ms, flip angle=72°, FOV=224×111 mm, matrix size=112×110 mm, number of slices=37, voxel size=2×2×2 mm3 (resting state scan). Note that the FOV for resting state scans was smaller compared to the task scans, which led some brain regions (i.e. predominantly in the visual cortex and cerebellum) to contain missing values when obtained during rest, but not during task performance. Each scan was carefully reviewed by a neuroradiologist, and any significant findings were reported to the subject’s primary care physician.

### fMRI preprocessing

Images were preprocessed using an in-house developed native space method.^39^ Briefly, slice timing correction is performed with FSL. slicetimer tool. We used mcflirt (motion correction tools in the FSL. package)^40^ to register all the volumes to a reference image (jenkinson et al,. 2002).^41^ The reference image was generated by registering (6 df, 256 bins mutual information, and Sinc interpolation) all volumes to the middle volume and averaging them. We then used the method described in Power et al. (2012) to calculate frame-wise displacement (FD) from the six motion parameters and root mean square difference (RMSD) of the BOLD percentage signal in the consecutive volumes for every subject, and used a threshold of .3%.^42^ RMSD was computed on the motion-corrected volumes before temporal filtering. The contaminated volumes were detected by the criteria FD>.5 mm or RMSD>.3%. Identified contaminated volumes were replaced with new volumes generated by linear interpolation of adjacent volumes. Volume replacement was done before band-pass filtering.^43^ The motion-corrected signals were passed through a band-pass filter with the cut-off frequencies of .01 and .09 Hz. We used flsmaths—bptf to do the filtering in this study.^40^ Finally, we residualized the motion-corrected, scrubbed, and temporally filtered volumes by regressing out the FD, RMSD, left and right hemisphere white matter, and lateral ventricular signals.^44^ Images that had undergone more than 30% scrubbing, were excluded from the dataset.

### Calculation of functional connectivity matrices

T1 image segmentation was done using FreeSurfer v5.1^45^ and visually checked for any inaccuracy. Corrections were made according to the Freesurfer provided guidelines (https://surfer.nmr.mgh.harvard.edu/fswiki/FsTutorial/TroubleshootingData). The coordinates of the 264 putative functional nodes, derived from a brain-wide graph that can be subdivided into multiple functional systems (e.g. default mode, visual, fronto-pańetal),^46^ was transferred to subjects T1 space with non-linear registration of the subjects structural scan to the MNI template using ANTS software package.^47^ A spherical mask with 10 mm radius and centered at each transferred coordinates was generated and intersected with the Freesurfer gray-matter mask to obtain the region of interest (ROI) mask for the 264 functional nodes. An intermodal, intrasubject, rigid-body registration of fMRI reference image and T1 scan was performed with FLIRT with 6 degree of freedom, normalized mutual information as the cost function,^48^ and used to transfer all the ROI masks from T1 space to FMRI space. These transferred ROI masks were used to average all the voxels within each mask to obtain a single FMRI time-series for each node. For each subject and each condition (rest, 11 RANN tasks), Pearson correlation coefficients were calculated for all possible pairs among the 264 time-series and Fisher z-transformed. We discarded four of these 264 nodes (three “uncertain” and one “default mode” region) for further analysis (and all connectivity pairs associated with them), as more than 20 subjects had missing resting state data in these regions.

### Calculation of task potency maps

Our approach to calculate task potency is similar (although not identical) to earlier papers on this method, of which a detailed explanation is provided elsewhere.^26,27^ In Matlab R2017a, we created vectors containing all unique pairs among the 260 × 260 connectivity matrices (i.e. after excluding the four nodes described above). We then standardized each task’s vector on a subject level by the vector created from resting state on a group level. More specifically, for each subject’s connectivity value in each pair, we subtracted the mean connectivity and divided by the standard deviation for that pair across all participants during resting state. This resulted in 11 “task potency” maps for every individual, reflecting pairwise changes in connectivity from the resting state to each task state. A positive task potency value reflects enhanced synchronicity between nodes during task performance, whereas negative task potency indicates reduced synchronicity (note that this could imply the occurrence of decoupling, but also an increased inverse coupling). Generally, a value that is further away from zero (irrespective of the direction) means a greater change in connectivity from resting state.

### Cortical thickness analysis

Using each individual’s T1-weighted MPRAGE image, cortical thickness measures were derived using the FreeSurfer v5.1 software package (http://surfer.nmr.mgh.harvard.edu/). Although the estimation procedure is automated, we manually checked the accuracy of the spatial registration and the white matter and gray matter segmentations following the analytic procedures outlined by Fjell and colleagues.^49^ Cortical thickness was measured by first reconstructing the gray/white matter boundary and the cortical surface,^50^ and the distances between these surfaces at each point across the cortical mantle were calculated. Using a validated automated labeling system,^51^ FreeSurfer divided the cortex into 68 different gyral-based parcellations, and calculated the mean thickness in each area. We used the global mean cortical thickness across these 68 areas in our analyses.

## Statistical analysis

### Demographic and clinical characteristics

We summarized demographic and clinical characteristics of our sample for different age groups, and additionally performed Pearson’s correlation analyses to determine the relationships between age, education, premorbid IQ, cognitive performance and global mean cortical thickness.

### Summarizing task potency across tasks within cognitive abilities

Since the RANN tasks were categorized into four latent cognitive abilities on a behavioral level (i.e. vocabulary, episodic memory, processing speed, and fluid reasoning), we examined whether the same clusters would also exist in our fMRI data. If the clusters identified on a neuroimaging level would be in accordance with the four latent cognitive abilities, this would provide face validity for the task potency method in general, and provide a rationale for data reduction by summarizing task potency across tasks belonging to the same cognitive ability. Therefore, we created 11 group average task potency maps, and determined the between-task correlation of potency values across pairs. In addition, we performed a k-means clustering analysis in Matlab (i.e. based on a group-level matrix with rows corresponding to tasks and columns to pairs) to create four clusters, based on the squared Euclidean distance, and a maximum of 100 iterations. We allowed four clustering repetitions with new initial cluster centroid positions, and used the solution with the lowest within-cluster sums of point-to-centroid distances. Based on the outcome (see results section) we created four new individual level potency maps for each cognitive ability.

### Identification of CR-related task-invariant connectivity pairs

To identify task-invariant networks related to CR, we performed linear regression analyses for all four cognitive abilities, with CR factors (i.e. either education or premorbid IQ) as predictors and task potency in each pair as the dependent variable. We selected all pairs in which the relationship between task potency and education or premorbid IQ was significant (p<.05) across cognitive abilities. Although we did not specify the direction of these relationships, we expected that within-pair relationships would be either consistently positive or negative in each cognitive ability. Furthermore, to account for multiple comparisons, we simulated our analyses with 1000 permutations in a random dataset (i.e. we randomly re-assigned the premorbid IQ and education scores among our subjects while maintaining their original task potency maps). With each permutation, we identified the number of pairs that showed significant relationships in the same direction with the CR proxies across cognitive abilities (Figure 2). This allowed us to compare the number of task-invariant pairs identified in our “real” dataset with the amount of significant pairs that would be observed by chance alone. To minimize false positive findings, we identified the threshold at which the number of significant pairs was below 5% in the distribution of the permuted data, and only considered education and IQ-related networks in which the number of pairs included were above this threshold for further analysis. To unburden the discussion of the results later on, we already report here that the thresholds for education and premorbid IQ were 20 and 19 pairs, respectively. Finally, we also determined which pairs showed a consistent relationship with age, since premorbid IQ was collinear with this variable (Table 2). This step enabled us to determine the degree of overlap between age- and CR-related pairs, as we wanted to rule out the possibility that the identified task-invariant CR networks actually resulted from an age effect.

**Figure 2.**
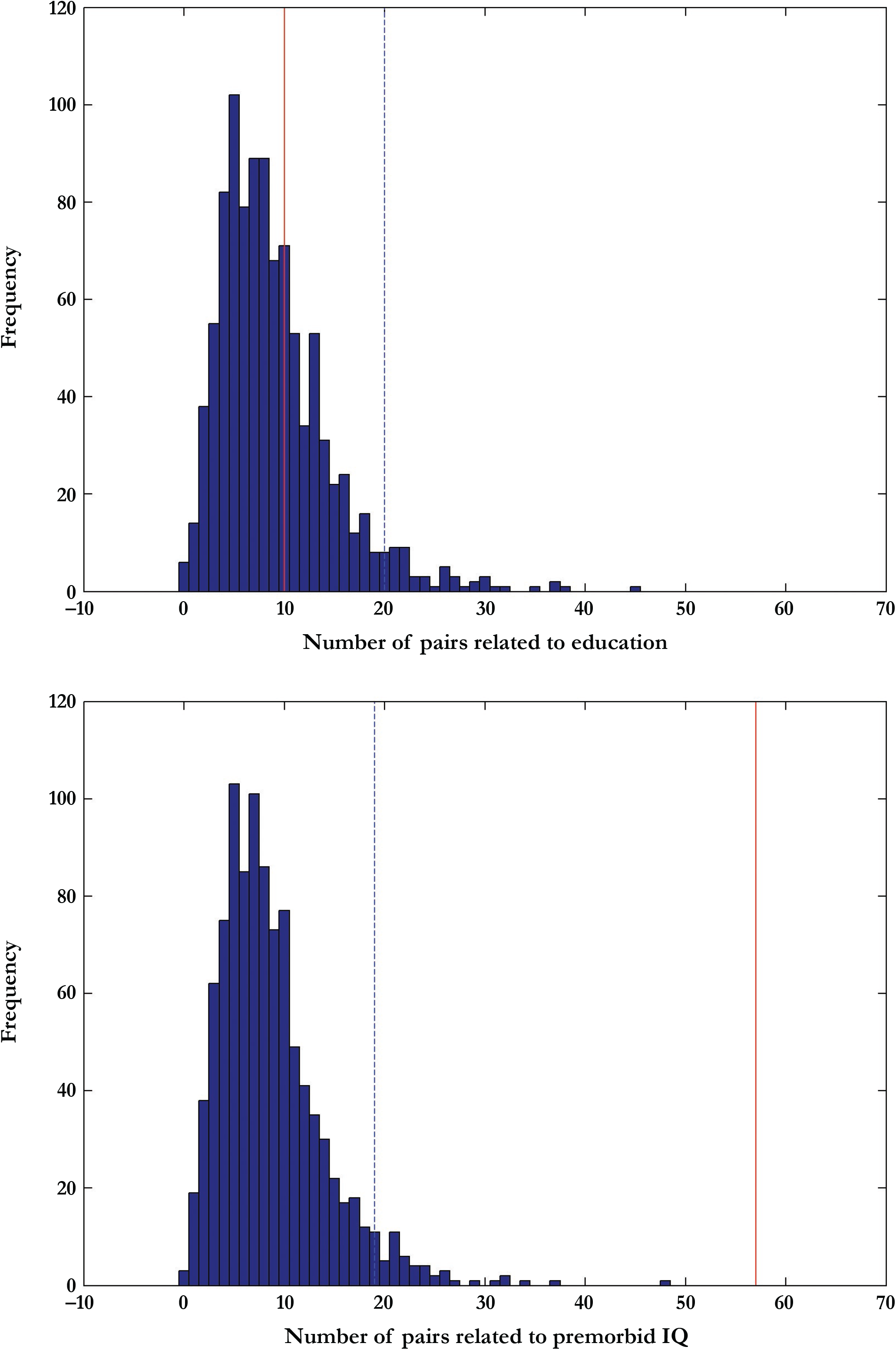
Distribution of the number of pairs that are significant across cognitive abilities in a random dataset. We performed 1000 permutations based on sampling without replacement. Red line=number of pairs found in actual dataset, dotted blue line=number of pairs found in random dataset.

**Table 1.** Characteristics of our sample according to different age groups.

| **Age** | **20-29** | **30-39** | **40-49** | **50-59** | **60-69** | **70-80** |
| --- | --- | --- | --- | --- | --- | --- |
| **N** | 42 | 47 | 39 | 44 | 49 | 44 |
| **Sex (% male)** | 31.0 | 31.9 | 56.4 | 54.5 | 55.1 | 45.5 |
| **Education (years)** | 15.64 ± 2.23 | 16.28 ± 2.39 | 16.13 ± 2.73 | 16.43 ± 1.82 | 16.16 ± 2.30 | 16.80 ± 2.76 |
| **IQ** | 113.55 ± 8.49 | 112.66 ± 8.94 | 113.72 ± 9.21 | 117.52 ± 7.51 | 119.78 ± 7.58 | 120.65 ± 7.16 |
| **DRS total** | 140.51 ± 2.51 | 139.89 ± 2.68 | 139.39 ± 2.68 | 140.32 ± 2.99 | 139.67 ± 3.13 | 139.64 ± 2.86 |
| **Global mean Cth** | 2.70 ± .11 | 2.65 ± .09 | 2.66 ± .09 | 2.61 ± .07 | 2.57 ± .10 | 2.53 ± .11 |

Data are displayed as mean ± SD. DRS=score on the Mattis Dementia Rating Scale, IQ=score on the American National Adult Reading Test, Cth=cortical thickness. Global mean cortical thickness was obtained by averaging across 68 Freesurfer parcellations.

**Table 2.** Relationships between age, CR factors, cortical thickness and cognitive performance.

|  | **Age** | **IQ** | **Education** | **VOCAB** | **MEM** | **SPEED** | **FLUID** | **Global mean Cth** |
| --- | --- | --- | --- | --- | --- | --- | --- | --- |
| **Age** | - | **.35\*\*** | .11 | **.35\*\*** | **-.34\*\*** | **.54\*\*** | **-.32\*\*** | **-.52\*\*** |
| **IQ** | **.35\*\*** | - | **.50\*\*** | **.74\*\*** | **.25\*\*** | .09 | **.31\*\*** | -.07 |
| **Education** | .11 | **.50\*\*** | - | **.37\*\*** | **.20\*** | -.03 | **.24\*\*** | -.01 |
| **VOCAB** | **.35\*\*** | **.74\*\*** | **.37\*\*** | - | **.27\*\*** | -.04 | **.29\*\*** | -.06 |
| **MEM** | **-.34\*\*** | **.25\*\*** | **.20\*** | **.27\*\*** | - | **-.41\*\*** | **.59\*\*** | **.24\*\*** |
| **SPEED** | **.54\*\*** | .09 | -.03 | -.04 | **-.41\*\*** | - | **.30\*\*** | **-.24\*\*** |
| **FLUID** | **-.32\*\*** | **.31\*\*** | **.24\*\*** | **.29\*\*** | **.59\*\*** | **.30\*\*** | - | **.29\*\*** |
| **Global mean Cth** | **-.52\*\*** | -.07 | -.01 | -.06 | **.24\*\*** | **-.24\*\*** | **.29\*\*** | - |

Results represent Pearson's correlation coefficients, \*\*=significant at $p < .001$ , \*=significant at $p < .01$ . IQ=score on the American National Adult Reading Test, Cth=cortical thickness. Global mean cortical thickness was obtained by averaging across 68 Freesurfer parcellations.

### Relationships between task potency and cognitive performance

To test whether the task-invariant CR network(s) identified in the previous step acted as a moderator between brain structure and cognition, we carried out linear regression analyses in which cognitive performance in either episodic memory, processing speed, and fluid reasoning was predicted from global mean cortical thickness (i.e. across 68 parcellations), task potency within the task-invariant CR network, and their interaction (Model 1). Vocabulary was not considered in this analysis, as task performance within this cognitive ability was uncorrelated with global mean cortical thickness (and thus we also did not expect a moderating effect of task potency). For cognitive abilities that showed no significant interaction between task potency and global mean cortical thickness, we also performed analyses without an interaction term (Model 2), examining the direct relationship between task potency and cognitive performance (i.e. after adjusting for brain structure). To summarize task potency across the CR-related pairs, we performed Principal Component Analysis (PCA) on centered task potency values and then used of each pair’s loadings to the first component (Supplementary Table 2) to create subject scores. These subject scores take into account the interdependence of task potency values across pairs, and treats task potency within the task-invariant CR network as one latent construct. A higher task potency summary score indicates that the network as a whole showed greater change (i.e. both positive and negative) from resting state connectivity, while a lower score reflects less change from resting state.

## Results

### Demographic and clinical characteristics

Table 1 provides demographic features of the study participants. As shown in Table 2, there was a positive correlation between age and premorbid IQ (r=.346, p<.001). Education and premorbid IQ were also positively related (r=.496, p<.001). Performance on each cognitive ability was related to scores within the other abilities (except for processing speed and vocabulary). Lower global mean cortical thickness was associated with older age (r=—.524, p<.001), confirming that this measure captures age-related changes in brain structure. Finally, higher age and lower global mean cortical thickness were related to worse cognitive performance (except for global mean cortical thickness and vocabulary).

### Summarizing task potency across tasks within cognitive abilities

We used group average task potency maps to determine the between-task correlation of potency values across pair. As shown in Figure 3, the correlations among within-ability tasks where generally higher than for between-ability tasks. This observation was confirmed by the k-means clustering analysis, which resulted in four clusters that were identical to the RANN latent cognitive abilities (Supplementary Figure 1). This provided a rationale for the summarization of task potency values within each cognitive ability, and we thus created four new task potency maps (i.e. for vocabulary, episodic memory, processing speed and fluid reasoning) for each individual. These maps were used for further analyses.

**Figure 3.**
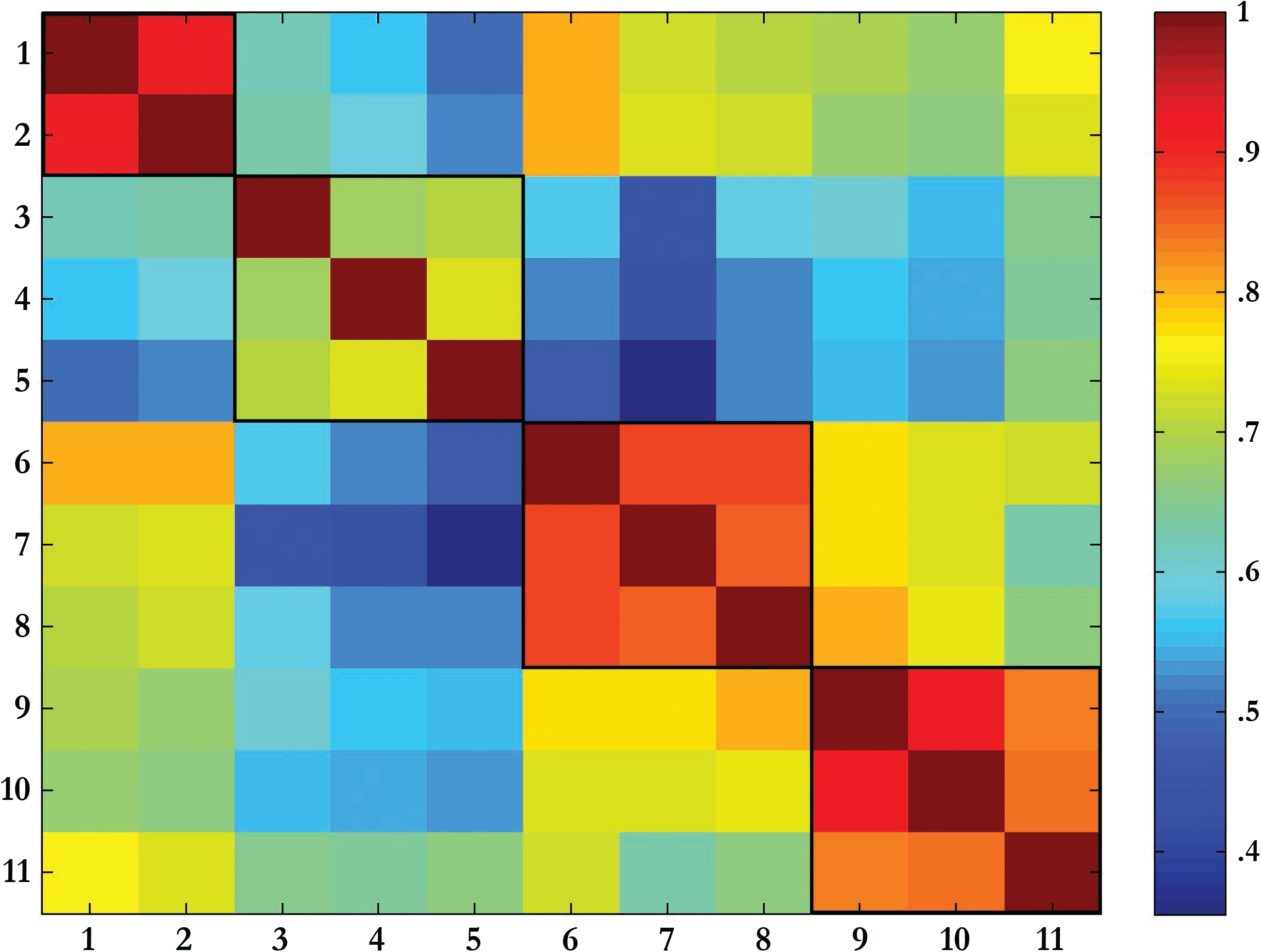
Correlation between group task potency maps for each RANN task. Tasks represented in ascending order: 1. Antonyms, 2. Synonyms (VOCAB); 3. Logical Memory, 4. Paired Associates, 5. Word Order Recognition (MEM); 6. Letter Comparison, 7. Pattern Comparison, 8. Digit Symbol (SPEED); 9. Matrix Reasoning, 10. Paper Folding, 11. Letter Sets (FLUID).

### Identification of CR-related task-invariant connectivity pairs

Linear regression analyses across all four cognitive abilities, revealed 10 pairs in which education was consistently related to task potency. As this number of pairs was below the threshold of 19, as established based on random permutations (see statistical analysis section), the probability that these pairs were observed by chance alone was high and thus we did not consider this network for further analysis. In contrast, we found 57 pairs in which premorbid IQ was significantly related to task potency across all cognitive abilities (Figure 4). The ROIs included in these pairs where mainly part of the default mode (21%), fronto-parietal task control (14%) and salience system (9%, see Supplementary Table 2). Among these pairs, there were both positive and negative correlations, but the direction of these relationships across cognitive abilities were always consistent within pairs. Finally, there was also a high number of pairs (i.e. 1317) in which task potency was task-invariantly related to age, but only 17 of these pairs overlapped with those associated with premorbid IQ. This means that the IQ-related task-invariant network was largely unique and could not be explained by an effect of age alone.

**Figure 4.**
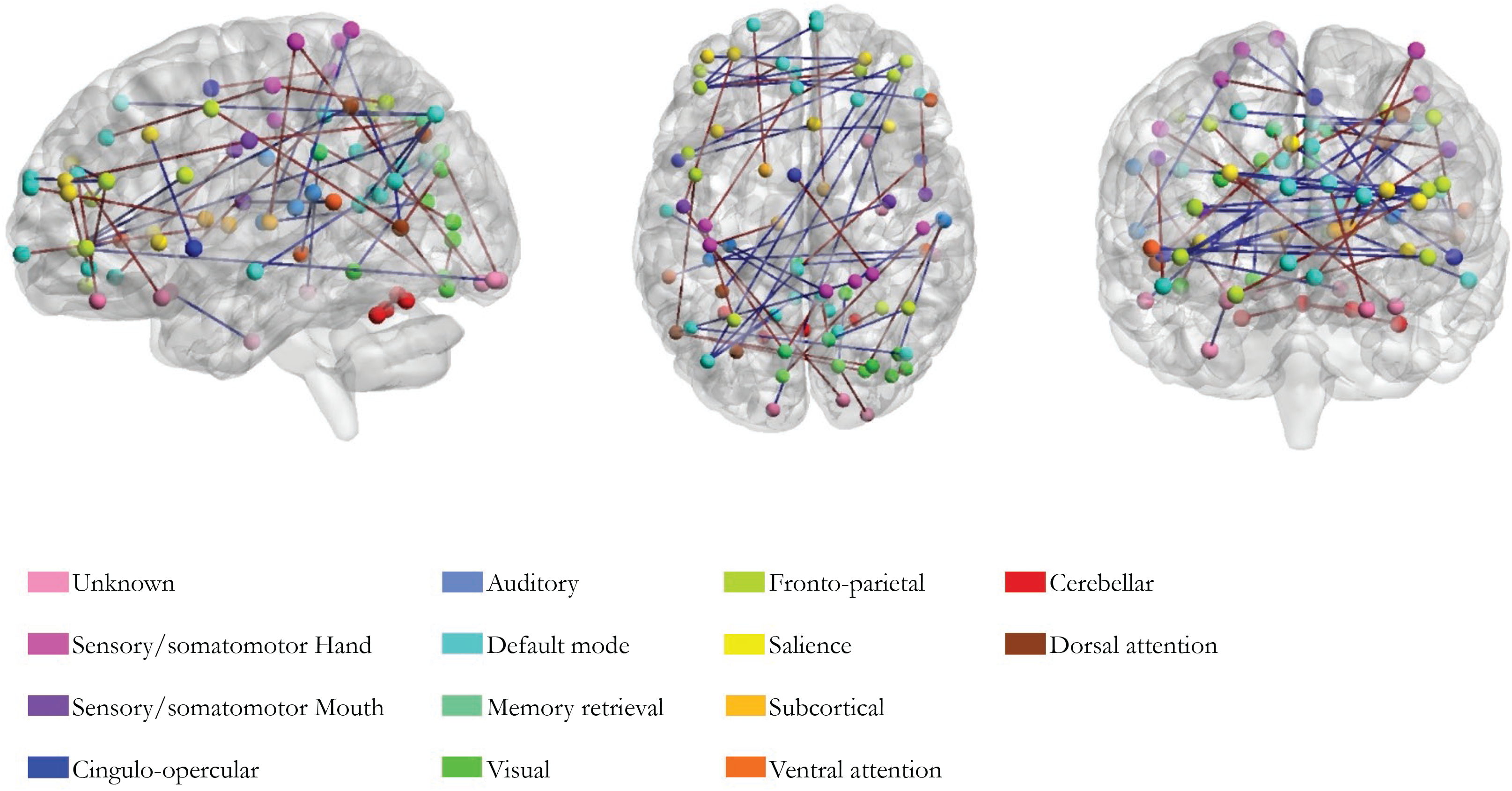
The IQ-related task-invariant network. ROIs with the same color belong to the same system as defined by Power et al (2011). Edges in blue reflect a negative relationship between premorbid IQ and that pair’s task potency, edges in red indicate a positive relationship. This figure was created with the BrainNet Viewer (http://www.nitrc.org/projects/bnv/) (Xia et al., 2013).

### Relationships between task potency and cognitive performance

The relationships of global mean cortical thickness, task potency in the IQ-related task-invariant network and their interaction with performance on the three cognitive abilities is summarized in Table 3. For episodic memory, we found main effects in Model 1 for both predictors (global mean cortical thickness: β=1.75, p<.001; task potency: β=1.58, p=.01) and an interaction effect (β =−.64, p=.01). Specifically, at higher levels of task potency (i.e. a greater change in connectivity relative to rest) within the IQ-related task-invariant network, the relationship between global mean cortical thickness and episodic memory performance was attenuated (Figure 5). For processing speed and fluid reasoning, there was only a main effect of global mean cortical thickness (β =−1.67, p<.001; β=2.43, p<.001, respectively – please note that the effect is in a opposite direction for processing speed because higher scores reflect slower reaction times and thus worse performance). Model 2, in which we assess the direct relationship of task potency with both processing speed and fluid reasoning while adjusting for the effects of global mean cortical thickness, showed a significant effect for fluid reasoning (β=.08, p<.01). This indicated that individuals with higher levels of task potency in the IQ-related task-invariant network performed better on this cognitive ability than subjects with a comparable brain structure but lower task potency. There was no direct relationship between task potency and processing speed (β =−.02, p<.38).

**Figure 5.**
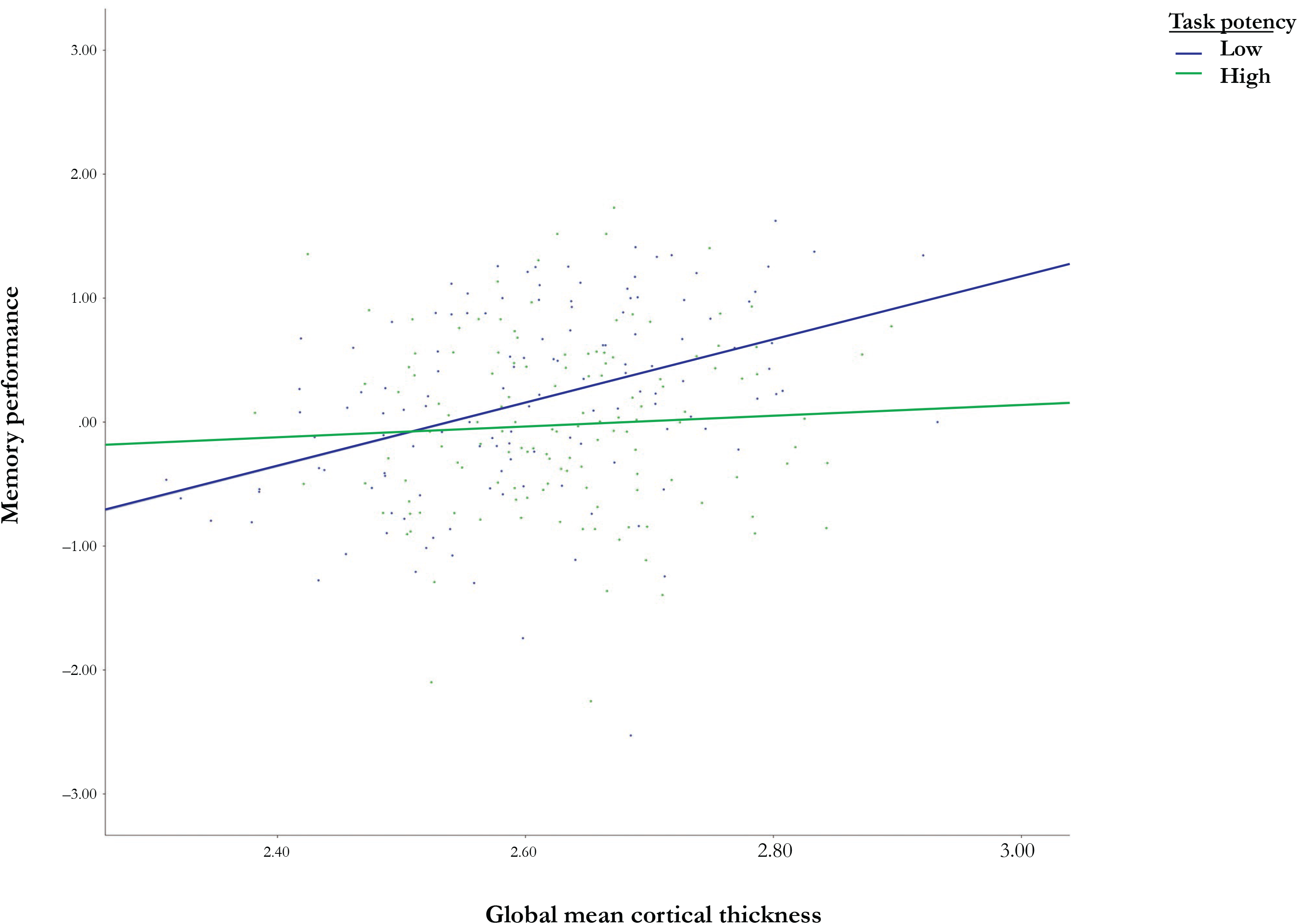
Moderation of the relationship between memory and cortical thickness by of task potency in IQ task-invariant network. We performed a median split of our sample into groups with low (mean=—1.28, SD=.86) and high task potency (mean=1.27, SD=1.19) within the IQ-related task-invariant network.

**Table 3.** The effects of global mean cortical thickness and task potency on cognitive performance.

|  | **MEM** |  | **SPEED** |  | **FLUID** |  |
| --- | --- | --- | --- | --- | --- | --- |
| **Model 1** | $\beta$ | **P** | $\beta$ | **P** | $\beta$ | **P** |
| **Global mean Cth** | **1.75\*\*** | **<.001** | **-1.67\*\*** | **<.001** | **2.43\*\*** | **<.001** |
| **TP** | **1.58** | **.01** | **-.28** | **.66** | **.47** | **.49** |
| **Global mean Cth\*TP** | **-.64** | **.01** | **.10** | **.69** | **-.15** | **.57** |
| **Model 2** |  |  |  |  |  |  |
| **Global mean Cth** | **-** | **-** | **-1.67\*\*** | **<.001** | **2.40\*\*** | **<.001** |
| **TP** | **-** | **-** | **-.02** | **.38** | **.08\*** | **<.01** |

$\beta$ =coefficient for the prediction of cognitive performance, Cth=cortical thickness, TP=task potency in the IQ-related task-invariant network. \*\*=significant at $p<.001$ , \*=significant at $p<.01$ .

## Discussion

### Summary of results

In this study, we used an adapted version of the task potency method^26,27^ to identify a task-invariant CR network in a large group of healthy subjects across the age span. This network consisted of 57 pairs of brain regions in which the change in functional connectivity during multiple cognitive tasks relative to rest (i.e. task potency), was directly related to premorbid IQ. The brain regions involved in this network were predominantly part of the default mode, frontoparietal task control and salience systems.^46^ Task potency in this network moderated the relationship between global mean cortical thickness and episodic memory, such that the impact of lower cortical thickness on memory performance was attenuated for individuals with greater task potency. In addition, after adjusting for cortical thickness, we found a direct effect of task potency on fluid reasoning: subjects with higher task potency performed better than persons with a comparable brain structure but lower task potency within the task-invariant CR network.

### Previous literature on task-invariant networks

We are not the first to investigate the existence of a task-invariant network underlying CR. In a sample of healthy subjects across the age span, Stern and colleagues identified a common spatial pattern of task load-related BOLD activity from the encoding phase of a working memory task that required two distinct cognitive processes: a verbal (i.e. letter) and object (i.e. shape) condition.^14^ This pattern showed correlations with premorbid IQ and vocabulary scores among young individuals, and involved brain regions in the superior and medial frontal gyrus.^14^ Using a more elaborate cognitive task battery, another study in a similar sample performed PCA on block and event-related contrasts of 12 RANN tasks (i.e. the same tasks we currently used, with the addition of Picture Naming) and found that the first principal component constituted a pattern that was common to all tasks, and also correlated with education. Positive loadings to this pattern were found within regions associated with the dorsal attention system, and negative loadings were within areas that were reminiscent of the default mode system.^12^ In a different study, Stern et al used the same RANN data and determined the number of principal components that optimally predicted premorbid IQ in a linear regression.^11^ Regression weights from this analysis were used to create an IQ-related task-invariant BOLD pattern, and expression of this pattern modulated between cortical thickness and fluid reasoning performance among healthy individuals across the age span. The most important areas that contributed to this pattern were the cerebellum, (superior) temporal, (inferior) parietal, precuneus and several regions within the frontal cortex.^11^ A study by Cole and colleauges among college students used a working memory task in which a verbal (i.e. words) and non-verbal (i.e. faces) was included.^15^ They showed that the lateral prefrontal cortex (LPFC) was involved in both conditions, and conceptualized the involvement of this area as related to cognitive control (i.e. activation levels correlated to the degree of control required, the correctness of the response and overall accuracy). Importantly, they subsequently used resting state fMRI to demonstrate that global connectivity of the LPFC was related to fluid intelligence.^15^ Further applying this finding to CR, Franzmeier and colleagues showed that resting state global connectivity of the LPFC correlated with education and also acted as a moderator in the relationship between hypometabolism in the precuneus and memory performance among individuals with prodromal Alzheimer’s disease.^16,17^

In our study, we combined favorable properties of both event-related and resting state fMRI techniques described above. That is, using an adapted version of the task potency method introduced by Chauvin and colleagues,^26,27^ we were able to acquire functional data during task performance, while concurrently taking into account the interconnected nature of the brain. The task potency measure is derived from an integration of both task-related and resting state functional connectivity. Functional connectivity during task performance likely reflects the sum of baseline connectivity and specific changes from this baseline in response to a given task. By extracting the difference between task-related and resting state functional connectivity, we arguably captured highly unique, task-related information. To our knowledge, we are the first to use task potency data to derive a task-invariant network in the context of CR. Interestingly, one study unrelated to CR recently demonstrated the existence of a common network in which task potency related to cognitive performance across three different fMRI tasks.^27^ Apart from visual and motor areas (which were expected to be involved due to the visual nature of these tasks and the motor responses that they required), the authors also found “higher-order” temporofrontal areas to be part of this network, which they attributed to the exertion of cognitive control across all tasks (i.e. a cognitive process that has previously been linked to CR).^16^

### Relevance of our findings

Our study supports the idea that CR operates, at least partly, in a task-invariant manner. This finding suggests that a single, generic mechanism contributes to the preservation of normal function across multiple cognitive abilities, which has two important implications. First, such a task-invariant network constitutes an ideal target for interventions studies. If we succeed at determining how to improve the performance of this network, it could protect individuals with initially low CR against the development many forms of cognitive decline with potentially various etiologies. Treatments aimed at enhancing CR have been suggested as a promising therapy to delay or prevent the emergence of cognitive decline.^52^ Secondly, the characterization of CR as a single mechanism on a functional brain level provides an alternative operationalization of the concept. Currently, CR is often measured based on proxies, which are easily measurable factors (e.g. education, premorbid IQ) that are correlated to the concept.^53^ However, proxies are often relatively static and therefore not suitable to capture within-individual variation in CR that results from the effects of aging and disease (i.e. decreases) or lifestyle enhancements (i.e. increases). Moreover, many proxies are not specific to CR, as they are also correlated with other resilience-related constructs, such as brain maintenance.^54^ Finally, proxies are conceptually unsatisfactory because they fail to distinguish between CR as a hypothetical construct and its determinants. Another increasingly popular approach to quantify CR is the use of residual,^55^ in which the discrepancy between brain structure and cognitive performance on an individual level is used as a more direct measure of the concept. While useful in a scientific context to elucidate the contributing factors and effects of CR, these methods provide a negative definition of the concept, which is not sufficient on a theoretical level. Although replication is needed, quantifying CR based on task potency in the task-invariant network we currently identified, could result in a more concrete, explanatory definition of CR.

### The absence of a task-invariant network for education

We identified a CR network that is task-invariant in the sense that task potency in the respective connectivity pairs correlated with premorbid IQ across all four cognitive abilities. In contrast, we did not find a task-invariant network for education. There are multiple explanations for this (absence of) result. First, it is possible that premorbid IQ, as compared to education, is more strongly related to CR.^56,57^ Indeed, it has been established that more intelligent individuals generally attain higher academic achievement,^58,59^ which suggests that (part of) the effect of education on CR could be explained by premorbid IQ. Moreover, education might be a less suitable measure of CR, because it has a more limited variance than premorbid IQ, especially in older cohorts and among women.^53,60^ It has also been argued that due to differences in ethnic backgrounds or socioeconomic status, academic quality is not uniform across individuals with equal years of education.^61^ Finally, heterogeneity in the type of education is not captured when this variable is measured in years. Individuals who complete many years of education tend to become increasingly specialized in a certain set of cognitive skills. For example, some persons may be particularly well-trained within the language domain, whereas other individuals with the same level of education might have further developed themselves on a mathematical level. As different cognitive abilities are (at least partly) coordinated by distinct, non-overlapping functional networks in the brain,^28^ education may be a less suitable candidate for the identification of a task-invariant network of CR. We speculate that education may impart CR through several independent mechanisms that support cognitive performance on a more task-specific level.^27^

### Task potency in the context of RANN

Our data were originally collected in the context of the RANN study, which aims to investigate whether spatial fMRI networks can be derived that are uniquely associated with the performance in each of the four cognitive abilities (i.e. vocabulary, episodic memory, processing speed, and fluid reasoning). Previous results have demonstrated convergent and discriminant validity for the 12 tasks included in the RANN study (i.e. all tasks that were used here, with the addition of Picture Naming) on a behavioral and functional brain level. Specifically, both cognitive performance scores and block-based BOLD activation patterns showed greater similarity between tasks within the same cognitive ability, and reduced similarity between tasks that reflected different cognitive abilities, respectively. In addition, linear indicator regression was used to derive four unique covariance patters for each cognitive ability, which showed good classification accuracy (i.e. identifying the correct task based on an individual’s fMRI activation pattern) in independent samples.^12,28^ Our results are in line with these reports, as we also found that the four cognitive abilities naturally emerged from our task potency data. That is, comparable to earlier findings, the correlations among group-level task potency maps from within-ability tasks where generally higher than for between-ability tasks, and using a k-means approach, four clusters could be identified that corresponded to each cognitive ability. These results have a two important implications. First, it further supports the existence of unique neural networks that underlie four main cognitive abilities that capture most of the age-related variance in cognitive performance. The fact that we replicated earlier findings with a novel fMRI analysis approach, suggests the robustness of these reference ability neural networks. Furthermore, the finding that our neuroimaging data could be summarized in a biologically meaningful manner and thus behaved as expected, provides additional face validity for the task potency method in general and demonstrates its utility and broad applicability.

### Strengths and limitations

This study has several strengths. We selected a relatively large, community-based sample of healthy individuals across a broad age range (i.e. 20-80). Importantly, each of these subjects underwent an elaborate fMRI procedure that included a resting state scan and 11 task-based scans. This within subjects-design is excellent for our research aim to a network that is truly involved in multiple tasks for each individual. Furthermore, we used a novel technique (i.e. task potency) to analyze the fMRI data, which provided unique, task-relevant information about the functional organization of the brain. One of the limitations of our study is that while most of the cognitive abilities were extensively assessed with three different tasks, our fMRI test batter only included two vocabulary tasks. It is therefore possible that the accuracy with which vocabulary performance was estimated, was somewhat lower compared to the other latent cognitive abilities. On a methodological level, another limitation is the fact that we excluded some subjects from our originally selected sample, as well as removed a set of ROIs from our analyses (i.e. due to large percentages of missing or scrubbed data). Excluded subjects, on average, were older and had a lower cortical thickness than the included sample. This implies that some degree of selection bias was at play, which may have affected the generalizability of our findings. Likewise, the exclusion of ROIs from our analysis may have caused us to overlook potentially relevant connectivity pairs in which task potency is (task-invariantly) related to CR. On a related note, for practical reasons, we decided to calculate task potency values based on group level resting state data, rather than using a subject level approach. There have been recent papers on the task potency technique that describe ways to obtain task potency values entirely based on subject-specific data.^26,27^ Briefly, this approach entails the subtraction of an individual’s resting state value in each connectivity pair from that subject’s task-based value, instead of standardizing it by the mean and standard deviation of the group average resting state data. This method accounts for possible differences between subjects in functional connectivity during rest, and is thus presumably more tailored towards the individual. On the other hand, without standardization to the group level, comparison between individuals becomes more difficult, which is a potential disadvantage of the subject-level task potency method in comparison to our current approach.

### Future studies

Our findings generate several areas for future study. First, a comparison between different task potency approaches (i.e. based on individual versus group level resting state data) is important to determine which provides the most meaningful data and thus should be the method of choice. Also, longitudinal data are useful to better understand how task potency in the task-invariant network specifically affects trajectories of decline in the four distinct cognitive abilities. This cross-sectional study revealed that task potency affected memory performance in a different manner (i.e. by moderating its association with brain structure) compared to fluid reasoning (i.e. through a direct positive relationship). These findings may suggest that the task-invariant CR network supports cognition in multiple ways, for example by affecting baseline performance (e.g. providing an initial advantage that is retained in the face of age- or disease-related structural brain changes) and causing longitudinal changes (e.g. allowing better preservation of premorbid cognitive function over time). In addition, since we showed that task potency was correlated with age in many connectivity pairs, it would also be interesting to examine how task potency itself changes over time as individuals age. Furthermore, cross-validation of our results in an independent sample or using different cognitive tests will be important to examine the robustness the task-invariant CR network and its effects on cognitive performance. Finally, other CR factors than education and premorbid IQ could be included in our approach. It would be informative to investigate whether occupation, for example, will also result in the identification of a task-invariant CR network, and to what degree it overlaps with the network we found based on premorbid IQ.

## Conclusion

In summary, we demonstrated that CR (at least partly) supports the ability to preserve cognitive function in a task-invariant manner. The identified task-invariant CR network, in which task potency related to premorbid IQ across four latent cognitive abilities, contributes to a better understanding of the mechanisms behind CR. This may in turn facilitate the development of new strategies to enhance CR and thereby minimize the negative impact of age- or disease-related structural brain changes on cognition. In addition, the task-invariant CR network could serve as a useful alternative operational measure of CR in a scientific context.

## Research highlights

– We measured task potency, which captures task-relevant changes in functional connectivity from rest, to identify a task-invariant network underlying CR among healthy individual across the age span.
–This network consisted of 57 connectivity pairs of regions of interest within the default mode, fronto-parietal and salience system, amongst others.
–Task potency in these pairs correlated with premorbid IQ (i.e. a proxy of CR) across four cognitive abilities: vocabulary, episodic memory, processing speed, and fluid reasoning.
–In addition, higher task potency within this task-invariant CR network attenuated the negative impact of lower cortical thickness on memory performance, and directly related to better fluid reasoning after adjusting for the effect of cortical thickness.

## Acknowledgements

We want to thank dr. R.J. Chauvin for her help and support in the implementation of the task potency method in our data. This project has been supported by the foundations “Alzheimer Nederland’ and “De Drie Lichten” in The Netherlands. These funding source(s) had no involvement in study design, collection, analysis and interpretation of data, or writing and submission of the report.

## References

1. Stern Y. Cognitive reserve in ageing and Alzheimer’s disease. The Lancet Neurology. 2012;11(11):1006–1012.

2. Arenaza-Urquijo EM, Wirth M, Chételat G. Cognitive reserve and lifestyle: moving towards preclinical Alzheimer’s disease. Front Aging Neurosci. 2015;7:134.

3. Bennett DA, Wilson RS, Schneider JA, Evans DA, Mendes de Leon CF, et al. Education modifies the relation of AD pathology to level of cognitive function in older persons. Neurology. 2003;60(12): 1909–1915.

4. Groot C, Hooghiemstra AM, Raijmakers PGHM, van Berckel BNM, Scheltens P, et al. The effect of physical activity on cognitive function in patients with dementia: a meta-analysis of randomized control trials. Ageing Res Rev. 2016;25:13–23.

5. Rentz DM, Huh TJ, Sardinha LM, Moran EK, Becker JA, et al. Intelligence quotient-adjusted memory impairment is associated with abnormal single photon emission computed tomography perfusion. J Int Neuropsychol Soc. 2007;13:821–831.

6. Scarmeas N, Zarahn E, Anderson KE, Habeck CG, Hilton J, et al. Association of life activities with cerebral blood flow in Alzheimer’s disease: implications for the cognitive reserve hypothesis. Arch Neurol. 2003;60:359–365.

7. Scarmeas N, Luchsinger JA, Schupf N, Brickman AM, Cosentino S, et al. Physical activity, diet, and risk of Alzheimer’s disease. JAMA. 2009;302:627–637.

8. Valenzuela MJ, Sachdev P. Brain reserve and cognitive decline: a non-parametric systematic review. Psychol Med. 2006;36:1065–1073.

9. Wilson RS, Barnes LL, Aggarwal NT, Boyle PA, Hebert LE, et al. Cognitive activity and the cognitive morbidity of Alzheimer’s disease. Neurology. 2010;75:990–996.

10. Wilson RS, Boyle PA, Yu L, Barnes LL, Schneider JA, Bennett DA. Life-span cognitive activity, neuropathologic burden, and cognitive aging. Neurology. 2013;81:314–321.

11. Stern Y, Habeck C. Deriving and testing the validity of cognitive reserve candidates. In Perneczky, RG (Ed.), Biomarkers for Preclinical Alzheimer’s Disease. 2018;63–70.

12. Habeck C, Gazes Y, Razlighi Q, Steffener J, Brickman A, et al. The Reference Ability Neural Network Study: life-time stability of reference-ability neural networks derived from task maps of young adults. Neuroimage. 2016;125:693–704.

13. Stern Y, Gazes Y, Razlighi Q, Steffener J, Habeck C. A task-invariant cognitive reserve network. Neuroimage. 2018;78:36–45.

14. Stern Y, Zarahn E, Habeck C, Holtzer R, Rakitin BC, et al. A common neural network for cognitive reserve in verbal and object working memory in young but not old. Cerebral Cortex. 2008;18(4):959–967.

15. Cole MW, Yarkoni T, Repovs G, Anticevic A, Braver TS. Global connectivity of prefrontal cortex predicts cognitive control and intelligence. J Neurosci. 2012;32(26):8988–8999.

16. Eranzmeier N, Caballero MÁA, Taylor ANW, Simon-Vermot L, Buerger K, et al. Resting-state global functional connectivity as a biomarker of cognitive reserve in mild cognitive impairment. Brain Imaging Behav. 2017;11(2):368–382.

17. Franzmeier N, Duering M, Weiner M, Dichgans M, Ewers M, Alzheimer’s Disease Neuroimaging Initiative (ADNI). Left frontal cortex connectivity underlies cognitive reserve in prodromal Alzheimer’s disease. Neurology. 2017;88(11):1054–1061.

18. van den Heuvel MP, Hulshoff Pol HE. Exploring the brain network: a review on resting-state fMKI functional connectivity. Eur Neuropsychopharmacol. 2010;20(8):519–534.

19. Hutchison RM, Womelsdorf T, Allen EA, Bandettini PA, Calhoun VD, et al. Dynamic functional connectivity: promise, issues, and interpretations. Neuroimage. 2013;80:360–378.

20. Gonzalez-Castillo J, Bandettini PA. Task-based dynamic functional connectivity: recent findings and open questions. Neuroimage. 2018;180(Pt B):526–533.

21. Braun U, Schäfer A, Walter H, Erk S, Romanczuk-Seiferth N, et al. Dynamic reconfiguration of frontal brain networks during executive cognition in humans. Proc Natl Acad Sci U S A. 2015;112(37): 11678–11683.

22. DeSalvo MN, Douw L, Takaya S, Liu H, Stufflebeam SM. Task-dependent reorganization of functional connectivity networks during visual semantic decision making. Brain Behav. 2014;4(6):877–885.

23. Fox MD, Snyder AZ, Zacks JM, Raichle ME. Coherent spontaneous activity accounts for trial-to-trial variability in human evoked brain responses. Nat Neurosci. 2006;9(1):23–25.

24. Smith SM, Fox PT, Miller KL, Glahn DC, Fox PM, et al. Correspondence of the brain’s functional architecture during activation and rest. Proc Natl Acad Sci U S A. 2009;106(31):13040–13045.

25. Tavor I, Parker Jones O, Mars RB, Smith SM, Behrens TE, Jbabdi S. Task-free MRI predicts individual differences in brain activity during task performance. Science. 2016;352(6282):216–220.

26. Chauvin RJ, Mennes M, Buitelaar JK, Beckmann CF. Assessing age-dependent multi-task functional co-activation changes using measures of task-potency. Dev Cogn Neurosci. 2018;33:5–16.

27. Chauvin RJ, Mennes M, Llera A, Buitelaar JK, Beckmann CF. Disentangling common from specific processing across tasks using task potency. NeuroImage. 2018;184:632–645.

28. Stern Y, Habeck C, Steffener J, Barulli D, Gazes Y, et al. The Reference Ability Neural Network Study: motivation, design, and initial feasibility analyses. Neuroimage. 2014;103:139–151.

29. Salthouse TA. Relations between cognitive abilities and measures of executive functioning. Neuropsychology. 2005;19:532–545.

30. Salthouse TA. Decomposing age correlations on neuropsychological and cognitive variables. J Int Neuropsychol Soc. 2009;15:650–661.

31. Salthouse TA, Pink JE, Tucker-Drob EM. Contextual analysis of fluid intelligence. Intelligence. 2008;36:464–486.

32. Mattis S. Dementia Rating Scale (DRS) Odessa, FL: Psychological Assessment Resources; 1988.

33. Blessed G, Tomlinson BE, Roth M. The association between quantitative measures of senile change in the cerebral grey matter of elderly subjects. British journal of Psychology. 1968;114:797–811.

34. Grober E, Sliwinski M. Development and validation of a model for estimating premorbid verbal intelligence in the elderly. J Clin Exp Neuropsychol. 1991;13:933–949.

35. Salthouse TA. Speed and knowledge as determinants of adult age differences in verbal tasks. J Gerontol. 1993;48:29–36.

36. Salthouse TA, Babcock RL. Decomposing adult age differences in working memory. Developmental Psychology. 1991;27:763–776.

37. Raven JC. Advanced progressive matrices, set 2. H.K. Lewis: London, UK; 1962.

38. Ekstrom RB, French JW, Harman HH, Dermen D. Manual for kit of factor-referenced cognitive tests. Educational testing service: Princeton, NJ; 1976.

39. Razlighi QR, Habeck C, Steffener J, Gazes Y, Zahodne LB, et al. Unilateral disruptions in the default network with aging in native space. Brain Behav. 2014;4(2):143–157.

40. Jenkinson M, Beckmann CF, Behrens TE, Woolrich MW, Smith SM. FSL. Neuroimage. 2012;62(2):782–789.

41. Jenkinson M, Bannister P, Brady M, Smith S. Improved optimization for the robust and accurate linear registration and motion correction of brain images. Neuroimage. 2002;17:825–841.

42. Power JD, Barnes KA, Snyder AZ, Schlaggar BL, Petersen SE. Spurious but systematic correlations in functional connectivity MRI networks arise from subject motion. Neuroimage. 2012;59(3):2142–2154.

43. Carp J. Optimizing the order of operations for movement scrubbing: comment on Power et al. Neuroimage. 2013;76:436–438.

44. Birn RM, Diamond JB, Smith MA, Bandettini PA. Separating respiratory-variation-related fluctuations from neuronal-activity-related fluctuations in fMRI. Neuroimage. 2006;31(4):1536–1548.

45. Fischl B. FreeSurfer. Neuroimage. 2012;62(2):774–781.

46. Power JD, Cohen AL, Nelson SM, Wig GS, Barnes KA, et al. Functional network organization of the human brain. Neuron. 2011;72(4):665–678.

47. Avants BB, Tustison N, Song G. Advanced normalization tools (ANTS). Insight J. 2009;2:1–35.

48. Jenkinson M, Smith S. A global optimisation method for robust affine registration of brain images. Med Image Anal. 2001;5(2): 143–156.

49. Fjell AM, Westlye LT, Amlien I, Espeseth T, Reinvang I, Raz N, Walhovd KB. High consistency of regional cortical thinning in aging across multiple samples. Cereb Cortex. 2009;19(9): 2001–2012.

50. Dale AM, Fischl B, Sereno MI. Cortical surface-based analysis. I. Segmentation and surface reconstruction. Neuroimage. 1999;9(2):179–194.

51. Fischl B, van der Kouwe A, Destrieux C, Halgren E, Ségonne F, et al. Automatically parcellating the human cerebral cortex. Cereb Cortex. 2004;14(1):11–22.

52. Stern Y. Cognitive reserve: implications for assessment and intervention. Folia Phoniatr Logop. 2013;65(2):49–54.

53. Jones RN, Manly J, Glymour MM, Rentz DM, Jefferson AL, Stern Y. Conceptual and measurement challenges in research on cognitive reserve, J Int Neuropsychol Soc. 2011;17(4):593–601.

54. Reed BR, Mungas D, Farias ST, Harvey D, Beckett L, et al. Measuring cognitive reserve based on the decomposition of episodic memory variance. Brain. 2010;133(8):2196–2209.

55. van Loenhoud AC, Wink AM, Groot C, Verfaillie SCJ, Twisk J, et al. A neuroimaging approach to capture cognitive reserve: application to Alzheimer’s disease. Hum Brain Mapp. 2017;38(9):4703–4715.

56. Jefferson AL, Gibbons LE, Rentz DM, Carvalho JO, Manly J, et al. Life course model of cognitive activities, socioeconomic status, education, reading ability, and cognition. J Am Geriatr Soc. 2011;59(8):1403–1411.

57. Richards M, Sacker A. Lifetime antecedents of cognitive reserve. J Clin Exp Neuropsychol. 2003;25:614–624.

58. Deary IJ, Strand S, Smith P, Fernandes C. Intelligence and educational achievement. Intelligence. 2007;35(1):13–21.

59. Lopes Soares D, Lemosa GC, Primi R, Almeida LS. The relationship between intelligence and academic achievement throughout middle school: the role of students’ prior academic performance. Learning and Individual Differences. 2015;41:73–78.

60. Starr J, Lonie J. Estimated pre-morbid IQ effects on cognitive and functional outcomes in Alzheimer’s disease: a longitudinal study in a treated cohort. 2008;8(1):27.

61. Manly JJ, Jacobs DM, Touradji P, Small SA, Stern Y. Reading level attenuates differences in neuropsychological test performance between African American and White elders. J Int Neuropsychol Soc. 2002;8(3):341–348.

