## Supplementary figures and images for "Identifying a task-invariant cognitive reserve network using task potency"

### Supplementary Figure 1

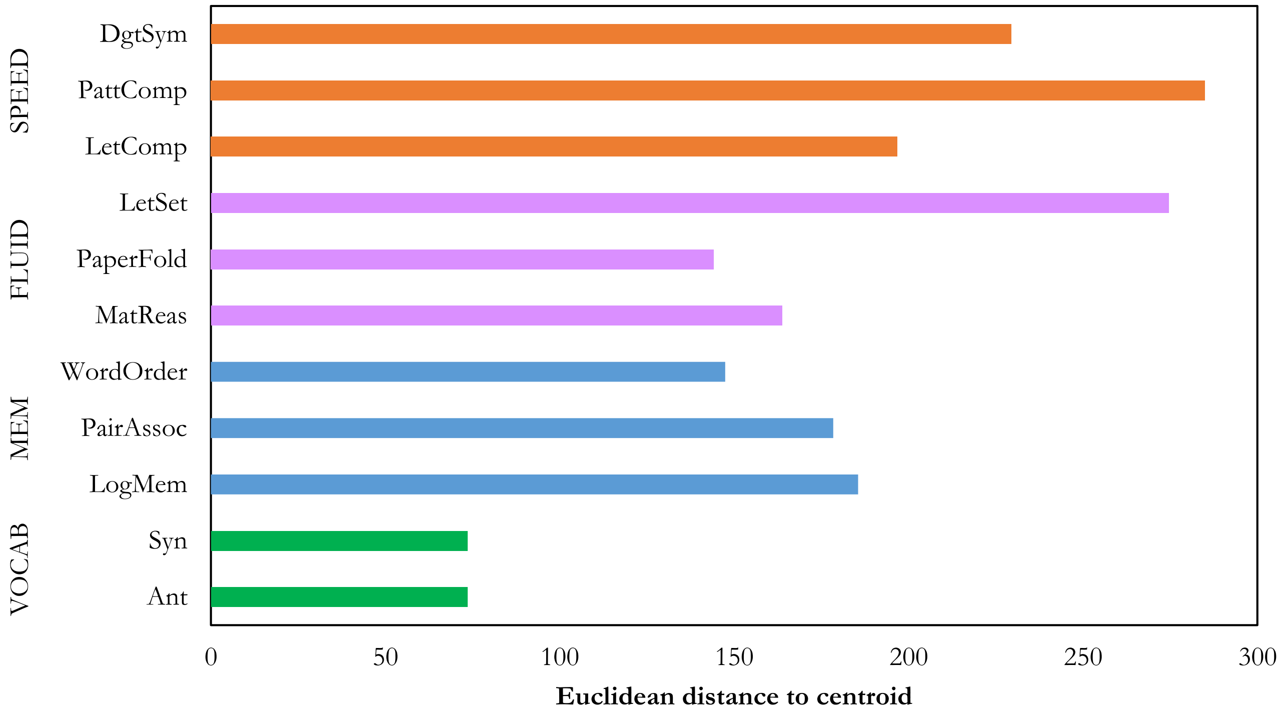
