## Supplementary tables for "Identifying a task-invariant cognitive reserve network using task potency"

**Supplementary Table 1. Characteristics of included and excluded RANN sample.**

|  | **Included (n=265)** | **Excluded (n=58)** | **p-value** |
| --- | --- | --- | --- |
| Age (range) | 50.04 (20-80) | 60.26 (24-77) | <.001 |
| Sex (%male) | 45.7 | 46.6 | .902 |
| Education (years) | 16.25±2.39 | 15.57±2.39 | .052 |
| DRS total | 139.90±2.83 | 140.02±3.01 | .778 |
| IQ | 116.53±8.67 | 115.45±9.19 | .407 |
| Mean global Cth | 2.618±.110 | 2.528±.128 | <.001 |

Data are displayed as mean±SD unless indicated otherwise. We used t-tests and Chi^2^ analysis (for sex) to compare both groups. DRS=score on the Mattis Dementia Rating Scale, IQ=score on the American National Adult Reading Test, Cth=cortical thickness.

**Supplementary Table 2. Overview of pairs that showed a significant relationship with IQ across all abilities.**

| **Network ROI 1** | **MNI (x, y, z)** | | | **Network ROI 2** | **MNI (x, y, z)** | | | **Relationship with IQ** |
| --- | --- | --- | --- | --- | --- | --- | --- | --- |
| Uncertain | 27 | –97 | –13 | Visual | 15 | –77 | 31 | + |
| Uncertain | 55 | –31 | –17 | Memory retrieval | –2 | –35 | 31 | – |
| Sensory/somatomotor Hand^*^ | 10 | –46 | 73 | Auditory^*^ | –30 | –27 | 12 | – |
| Sensory/somatomotor Hand | –40 | –19 | 54 | Default mode | –2 | 38 | 36 | + |
| Sensory/somatomotor Hand^*^ | –40 | –19 | 54 | Fronto–parietal Task Control^*^ | –28 | –58 | 48 | + |
| Sensory/somatomotor Hand | 29 | –39 | 59 | Cingulo–opercular Task Control | –3 | 2 | 53 | + |
| Sensory/somatomotor Hand | 50 | –20 | 42 | Visual | –8 | –81 | 7 | + |
| Sensory/somatomotor Hand^*^ | –38 | –27 | 69 | Uncertain^*^ | 17 | –91 | –14 | + |
| Sensory/somatomotor Hand^*^ | –38 | –27 | 69 | Subcortical^*^ | –10 | –18 | 7 | + |
| Sensory/somatomotor Hand^*^ | 22 | –42 | 69 | Auditory^*^ | 58 | –16 | 7 | – |
| Sensory/somatomotor Hand^*^ | 22 | –42 | 69 | Default mode^*^ | –46 | –61 | 21 | – |
| Sensory/somatomotor Mouth | –49 | –11 | 35 | Memory retrieval | –7 | –71 | 42 | + |
| Sensory/somatomotor Mouth | –49 | –11 | 35 | Salience | –28 | 52 | 21 | + |
| Sensory/somatomotor Mouth^*^ | 36 | –9 | 14 | Ventral attention^*^ | –55 | –40 | 14 | – |
| Sensory/somatomotor Mouth | 51 | –6 | 32 | Default mode | 49 | 35 | –12 | + |
| Cingulo–opercular Task Control | 49 | 8 | –1 | Default mode | –2 | –37 | 44 | + |
| Cingulo–opercular Task Control | –51 | 8 | –2 | Salience | 5 | 23 | 37 | – |
| Auditory | 58 | –16 | 7 | Visual | 40 | –72 | 14 | – |
| Auditory | –38 | –33 | 17 | Ventral attention | 51 | –29 | –4 | – |
| Auditory | 59 | –17 | 29 | Visual | 26 | –79 | –16 | + |
| Default mode | 6 | 67 | –4 | Subcortical | 9 | –4 | 6 | + |
| Default mode | –46 | –61 | 21 | Default mode | 43 | –72 | 28 | – |
| Default mode | –46 | –61 | 21 | Dorsal attention | –52 | –63 | 5 | – |
| Uncertain^*^ | 27 | 16 | –17 | Uncertain^*^ | 33 | –12 | –34 | – |
| Default mode | –39 | –75 | 44 | Default mode | –11 | –56 | 16 | – |
| Default mode | –39 | –75 | 44 | Default mode | –3 | –49 | 13 | – |
| Default mode | –39 | –75 | 44 | Default mode | –2 | –37 | 44 | – |
| Default mode^*^ | –39 | –75 | 44 | Default mode^*^ | 23 | 33 | 48 | – |
| Default mode | 8 | –48 | 31 | Memory retrieval | –7 | –71 | 42 | + |
| Default mode | 8 | –48 | 31 | Memory retrieval | 11 | –66 | 42 | + |
| Default mode | 15 | –63 | 26 | Fronto–parietal Task Control | 33 | –53 | 44 | + |
| Default mode | 11 | –54 | 17 | Default mode | –56 | –13 | –10 | – |
| Default mode | 6 | 64 | 22 | Fronto–parietal Task Control | –42 | 38 | 21 | – |
| Default mode | –3 | 44 | –9 | Salience | –28 | 52 | 21 | + |
| Default mode | –20 | 64 | 19 | Subcortical | –15 | 4 | 8 | + |
| Default mode^*^ | –8 | 48 | 23 | Uncertain^*^ | –21 | 41 | –20 | + |
| Uncertain | –31 | 19 | –19 | Salience | 26 | 50 | 27 | + |
| Memory retrieval | 11 | –66 | 42 | Fronto–parietal Task Control | 44 | –53 | 47 | – |
| Uncertain | –12 | –95 | –13 | Fronto–parietal Task Control | 43 | 49 | –2 | – |
| Visual^*^ | 18 | –47 | –10 | Visual^*^ | 15 | –77 | 31 | – |
| Visual | 43 | –78 | –12 | Visual | 37 | –81 | 1 | + |
| Visual^*^ | 29 | –77 | 25 | Dorsal attention^*^ | –52 | –63 | 5 | + |
| Visual | 26 | –79 | –16 | Dorsal attention | –52 | –63 | 5 | + |
| Fronto–parietal Task Control | –44 | 2 | 46 | Dorsal attention | –52 | –63 | 5 | + |
| Fronto–parietal Task Control^*^ | –47 | 11 | 23 | Fronto–parietal Task Control^*^ | 43 | 49 | –2 | – |
| Fronto–parietal Task Control | 24 | 45 | –15 | Fronto–parietal Task Control | –42 | 38 | 21 | + |
| Fronto–parietal Task Control | –42 | 38 | 21 | Fronto–parietal Task Control | 38 | 43 | 15 | – |
| Fronto–parietal Task Control | –42 | 38 | 21 | Salience | –28 | 52 | 21 | – |
| Fronto–parietal Task Control | –42 | 38 | 21 | Salience | 26 | 50 | 27 | – |
| Fronto–parietal Task Control | –42 | 45 | –2 | Ventral attention | 53 | 33 | 1 | – |
| Fronto–parietal Task Control | 43 | 49 | –2 | Dorsal attention | –33 | –46 | 47 | – |
| Fronto–parietal Task Control | 43 | 49 | –2 | Dorsal attention | –27 | –71 | 37 | – |
| Salience^*^ | –35 | 20 | 0 | Salience^*^ | 36 | 22 | 3 | – |
| Salience | 26 | 50 | 27 | Salience | –39 | 51 | 17 | – |
| Cerebellar^*^ | –16 | –65 | –20 | Cerebellar^*^ | 1 | –62 | –18 | + |
| Cerebellar^*^ | –32 | –55 | –25 | Cerebellar^*^ | 1 | –62 | –18 | + |
| Cerebellar^*^ | 22 | –58 | –23 | Cerebellar^*^ | 1 | –62 | –18 | + |

ROI=region of interest and its system membership according to Power,^46^ relationship IQ: + a positive relationship across all abilities between task potency and IQ, – a negative relationship between task potency and IQ, ^*^pairs in which task potency is also related to age.
